# *DUSP6* loss enhances dopamine secretion and modulate neurodevelopment in ADHD iPSC-derived dopaminergic neurons

**DOI:** 10.1101/2024.11.18.624053

**Authors:** Atefeh Namipashaki, Shane D. Hellyer, Aurina Arnatkeviciute, Cameron J Nowell, Kevin Walsh, Jose M. Polo, Karen J Gregory, Mark A. Bellgrove, Ziarih Hawi

## Abstract

Although the gene *DUSP6* has been implicated as a risk gene for attention deficit hyperactivity disorder (ADHD) in recent GWAS studies, its functional role in the aetiology of the condition remains poorly understood. *DUSP6* is reported to regulate dopaminergic neurotransmission by decreasing available synaptic dopamine, suggesting a potential mechanism by which *DUSP6* may confer risk to ADHD. In this study, we employed CRISPR–Cas9 to generate isogenic iPSC lines from an individual with ADHD, including heterozygous and homozygous *DUSP6* knockouts alongside the parental control. These iPSC lines were differentiated into dopaminergic neurons and assessed at both cellular, and transcriptomic levels. We observed significant increase in extracellular dopamine in proportion to *DUSP6* gene dosage, with greater elevation in homozygous compared to heterozygous knockouts. RNA sequencing further supported this finding, showing that *DUSP6*-knockout increased expression of genes regulating dopamine secretion and synaptic processes critical for dopaminergic system. Beyond dopamine signalling, transcriptomic changes implicated *DUSP6* in neurodevelopmental biological mechanisms, essential to the development of ADHD. Furthermore, we identified significant overlaps between *DUSP6*-regulated genes and those associated with major depression, bipolar disorder, and schizophrenia, suggesting shared genetic pathways potentially influenced by *DUSP6*. This study provides mechanistic insights into the role that *DUSP6* could potentially play in the aetiology of ADHD and its broader implications across related neuropsychiatric conditions.

## Introduction

Dual Specificity Phosphatase 6 (*DUSP6*), also known as *MKP3*, has been identified as a risk locus for ADHD in the recent large-scale genome-wide association studies (GWAS) (Demontis et al., 2023; Demontis et al., 2019; Faraone et al., 2024). However, the molecular mechanisms by which *DUSP6* contributes to ADHD risk remain incompletely understood.

*DUSP6* encodes a dual-specificity phosphatase that dephosphorylates both phosphoserine/threonine and phosphotyrosine residues (Seternes et al., 2019). Through regulation of kinase activity, *DUSP6* regulates diverse cellular processes including the regulation of dopaminergic neurotransmission. This is particularly relevant since dysregulation of dopamine has long been hypothesised as central to the aetiology of ADHD, with psychostimulant medications working by modulating dopamine signalling and increasing synaptic dopamine availability (Faraone et al., 2024). Previous research has shown that *DUSP6* can reduce synaptic dopamine levels (Mortensen, 2013; Mortensen et al., 2008), in part via downregulation of L-type voltage-gated calcium channel *CACNA1C*, which inhibits dopamine release (Mortensen, 2013), and stabilization of primary dopamine transporter (SLC6A3, previously known as DAT1) which increases dopamine reuptake (Mortensen et al., 2008). These findings suggest *DUSP6* may act as a risk gene by reducing synaptic dopamine levels, which is hypothesised as central to the aetiology of ADHD.

We therefore hypothesized that downregulating *DUSP6* in dopaminergic neurons derived from individuals with ADHD would be associated with increased synaptic dopamine levels. Using CRISPR–Cas9, we generated isogenic *DUSP6* heterozygous (Het) and homozygous (Hom) knockout iPSC lines and differentiated them into dopaminergic neurons. We quantified intra-and extracellular dopamine concentrations and assessed transcriptomic changes across the isogenic lines. Given that *DUSP6* overexpression reportedly downregulates the calcium channel *CACNA1C*, calcium imaging was performed to examine whether *DUSP6* knockout neurons exhibited enhanced calcium influx. Our findings suggest gene-dosage effect of the absence of *DUSP6*, leading to enhanced synaptic dopamine levels, thus offering potential mechanistic insights into the pathophysiology of ADHD.

## Materials and Method

### iPSC culture

An iPSC line derived from peripheral blood mononuclear cells (PBMC) of a male youth with ADHD was used (Tong et al., 2019). Cells were cultured on Biolaminin-521 (Biolamina, Cat#LN521) in StemFlex medium (Gibco, Cat#A3349401).

### CRISPR guide RNA (gRNA) design

The Alt-R CRISPR–Cas9 CRISPR RNA (crRNA) selection tool from Integrated DNA Technology (IDT) (https://sg.idtdna.com/site/order/designtool/index/CRISPR_PREDESIGN) was used to design the gRNA targeting exon 1 of the *DUSP6* gene. gRNA and primer sequences for the target and top two predicted off-target sites are listed in Supplementary Table 1.

### CRISPR mediated gene editing

CRISPR engineering of the iPSC line was conducted using our previously published protocol, which enables the generation of homogeneous edited iPSC clones within two weeks (Namipashaki et al., 2023). Briefly, equimolar (1 μM) gRNA oligonucleotides and Cas9 enzyme were combined to assemble the ribonucleoprotein complex. Dissociated iPSC clusters (150,000 cells) were transfected with the ribonucleoprotein and Lipofectamine Stem Transfection Reagent (Invitrogen, Cat#STEM00015) and seeded in 24-well plate. After 48 hours, positively transfected cells were sorted and plated as single cells in 96-well plates. Clones were cultured for 10 days to allow colony formation and screened by Sanger sequencing to identify correctly edited lines.

### Western blotting

Western blotting was used to confirm DUSP6 Het and Hom knockouts. Proteins were extracted from iPSCs using M-PER reagent (Thermo, #78501) with protease/phosphatase inhibitors (#78440), incubated on ice for 15 min, and centrifuged at 15,000 g for 15 min at 4 °C. Supernatants were quantified using the Qubit Protein Assay Kit (#A50668), separated on NuPAGE Bis-Tris gels (Invitrogen, #NP0335BOX), and transferred to PVDF membranes (#IB24002). Membranes were blocked (5% milk), incubated overnight with primary anti-DUSP6 antibody (Abcam, #ab76310), and incubated with HRP-conjugated secondary antibody. Detection used ECL Prime (Cytiva, #RPN2232) and ChemiDoc imaging (Bio-Rad).

### Dopaminergic neuron differentiation

The isogenic lines were differentiated into dopaminergic neurons in three independent replicate using a lentiviral induction method (Powell et al., 2023). iPSCs were transduced with a lentiviral system inducibly expressing ASCL1, LMX1B, and NURR1 along with PuroR. Cells were cultured on Biolaminin-111 LN in DMEM/F12 medium (Gibco, #10565018) with doxycycline (Sigma, #D9891) to activate transgene expression and puromycin (Sigma, #7255) for selection. At day 14 (DIV14), medium was switched to BrainPhys (Stem Cell Technologies, #05790) with growth factors and maintained until neuronal maturation at DIV35.

### Immunocytochemistry

To confirm dopaminergic identity, the developed neurons were assessed for the expression of tyrosine hydroxylase (TH); the rate limiting enzyme in catecholamine biosynthesis and key dopaminergic marker, and microtubule associated protein 2 (MAP2) neuronal marker. On day 35, cultures were washed with PBS, fixed in 4% paraformaldehyde for 15 min, permeabilized with 0.5% Triton X-100, and blocked in 3% BSA for 30 min. Cells were incubated with primary antibodies, anti–TH (Abcam, #ab75875) and anti–MAP2 (Abcam, #ab92434), for 2 hours at 37 °C, followed by Alexa Fluor secondary antibodies (Abcam, #ab150081, #ab150174) for 1 hour. NucBlue stain (Thermo, #R37605) was applied before imaging on an EVOS M5000.

### RNA isolation, library preparation, and sequencing

RNA was extracted using TRIzol and purified with the RNeasy Kit (Qiagen, #74104). Quantification, QC, library preparation, and sequencing were conducted at the Australian Genome Research Facility (AGRF). After confirming RNA quantity and integrity (Supplementary Table 2), 100 ng of RNA per sample was processed using the Illumina Ribo-Zero Plus workflow, which includes rRNA depletion, fragmentation (∼150 bp), and cDNA synthesis. Libraries were indexed, PCR-amplified, and sequenced on the Illumina NovaSeq X Plus platform at a depth of ∼50 million reads per sample.

### RNA-seq data processing and differential gene expression and enrichment analysis

Paired-end RNA-seq reads were processed using the nf-core/rnaseq pipeline (Harshil Patel, 2023). Adapters and low-quality reads were trimmed with Trim Galore v0.6.7 (Felix Krueger, 2021), and sequences were aligned to the GRCh38 human genome using STAR with ENSEMBL v109 annotations (Dobin et al., 2013). Aligned BAM files were converted to FASTQ using Samtools v1.14 and merged with unmapped reads (Patro et al., 2017). Gene-level counts normalized for transcript length were generated using tximport (Soneson et al., 2015). Differential expression analysis was performed in Degust using the limma-voom method (Law et al., 2014; Powell., 2019). Differentially expressed genes (DEGs) with a false discovery rate (FDR) < 0.05 were visualized in volcano plots for each pairwise comparison between parental, Het, and Hom lines. Top 10 genes from each comparison were ranked based on significance. The list of DEGs that exhibited consistent directional changes across Het, and Hom lines compared to the parental line was selected to represent the DEGs altered due to the gene–dosage effect of *DUSP6*, referred to as *DUSP6* dosage-responsive DEGs. These, in addition to up- and downregulated DEGs (FDR < 0.05) from each pairwise comparison underwent Gene Ontology (GO) enrichment analysis using the STRING database (Law et al., 2014; Szklarczyk et al., 2023). Enrichment strength was expressed as Log_10_ (observed/expected), and statistical significance was adjusted using the Benjamini-Hochberg method to control the false discovery rate (Szklarczyk et al., 2021).

### Analysis of *DUSP6*–DEG overlap with GWAS identified genes for major psychiatric disorders

Given that *DUSP6* has also been implicated as a risk locus for other psychiatric disorders, we examined the overlap between DEGs from each pairwise comparison (parental, Het, Hom) and GWAS-identified genes for major psychiatric disorders, including schizophrenia (SCZ), major depression (MDD), bipolar (BD), and autism spectrum disorder (ASD). Risk genes were derived using the MAGMA software (de Leeuw et al., 2015), mapping SNPs to genes with 1000 Genomes (European ancestry) LD reference files from https://github.com/thewonlab/H-MAGMA. Gene-level associations were computed using multiple linear principal component regression, with F-tests applied to generate p-values. Bonferroni correction (p-value < 0.05/total genes) identified significant loci as: 95 (ADHD), 336 (BD), 465 (MDD), 922 (SCZ), and 14 (ASD) genes. Overlap significance between ADHD–DEGs and GWAS–identified genes for major psychiatric conditions was evaluated using 100,000 permutations, comparing observed overlaps to randomly sampled gene lists of equal size. Overlapped genes for each disorder were subsequently analysed for their overrepresentation in biological processes using The Gene Ontology resource (de Leeuw et al., 2015).

### Dopamine ELISA measurement of intra- and extracellular dopamine

Intra and extracellular dopamine concentrations from three independent replicates were quantified using the Dopamine Research ELISA Kit (ALPCO, #17**-**DOPHU**-**E01**-**RES). Media supernatants were first collected and flash**-**frozen in liquid nitrogen. The neurons were then imaged using the cell viability stain (Thermo# A15001) to calculate the area of viable cells. Subsequently, they were harvested using Accutase and spun at 1000 × g for 5 minutes. Cell pellets were lysed by sonication and subjected to ELISA according to the manufacturer’s instructions. Absorbance at 450 nm was measured using a CLARIOstar Plus Microplate Reader and dopamine concentrations were normalized to micromoles per relative fluorescence unit (µMols/RFU) of viable cells. A linear trend across Parental, Het and Hom lines was assessed using One-Way ANOVA with Polynomial linear contrast, followed by pairwise comparisons (Parental vs Hom; Parental vs Het; Het vs Hom). Bonferroni corrections were applied to control for multiple comparisons (adjusted significance threshold p-value < 0.05/3).

### Calcium imaging

At DIV42 of dopaminergic differentiation, neurons underwent two half-media changes with calcium assay buffer, were loaded with Fluo-8 dye (1.6 µM) for 45 min at 37 °C and subsequently subjected to two half-volume washes. Imaging was performed using an Instant Structured Illumination Microscope (ISIM; VisiView v5.0.0.27) with a 100× oil objective (UPLAPO100XOHR, 1.50NA) and 488 nm excitation, captured via a Prime BSI sCMOS camera (65 nm pixel size, 1470 × 986 px, 20 ms exposure). Calcium imaging was conducted on multiple wells of growing neurons recorded at least 80 neurons for each genotype (parental, Het, and Hom) over a 90 s period, comprising 45 s baseline and 43.5 s response to 100 mM KCl stimulation. The entire recording period was divided into epochs: baseline (-35 to -25 s), pre-stimulus (-25 to -5 s), stimulus onset (-5 to 8 s), and stimulus response (8 to 20 s). Data were normalized to baseline (-35 to -25 s) and expressed as ΔF/F_0_, where F_0_ is mean baseline fluorescence. Statistical analysis was performed using mixed-effects modelling in MATLAB.

## Results

### CRISPR engineering and functional validation of heterozygous and homozygous *DUSP6* knockout dopaminergic neuron models

Using our previously published CRISPR engineering method (Namipashaki et al., 2023) which efficiently generates homogeneous single-cell CRISPR-edited clones, we successfully generated two *DUSP6* knockout clones with frameshift mutations. The first showed single-allele disruption (Het), while the second had biallelic disruption (Hom), confirmed by Sanger sequencing (Figure 1A). In silico analyses using the PHYRE2 protein prediction tool (Kelley et al., 2015) verified the respective disruptions and revealed altered protein conformations in both clones (Figure 1B). Western blotting further confirmed the absence of DUSP6 protein in the knockout lines, validating the successful generation of gene-dosage defined isogenic lines (Parental, Het, and Hom) that differ only at a single genomic locus (Figure 1C).

**Figure 1.**
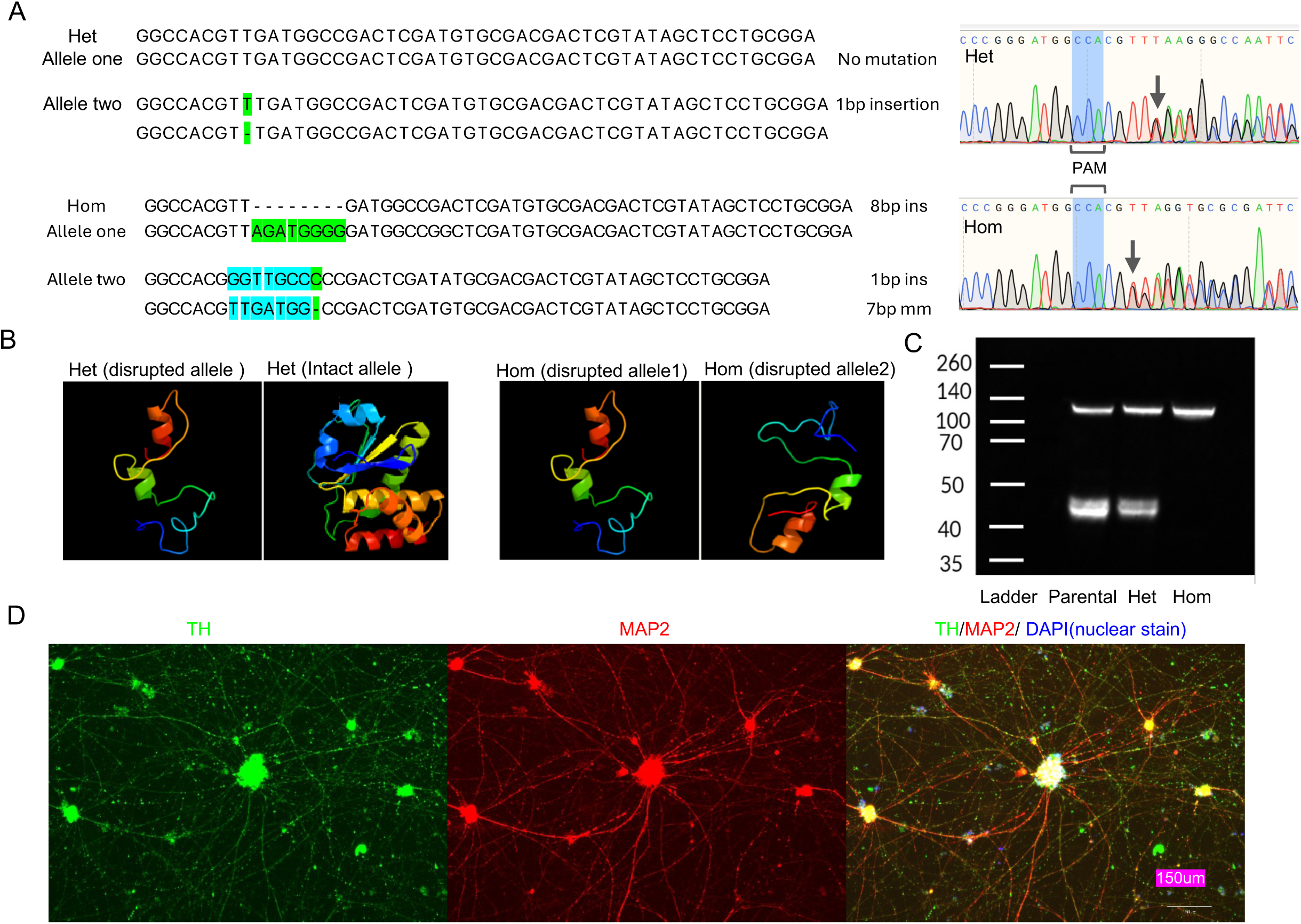
Generation of isogenic *DUSP6* knockout neuronal models. **A)** CRISPR–Cas9 editing generated two homogeneous single-cell iPSC clones: one Het and one Hom, as confirmed by Sanger sequencing. The Het line is characterized by a 1-base insertion in one allele, while the Hom line is characterized by a 1-base insertion in one allele and an 8-base insertion in the other allele of *DUSP6* exon one. **B)** PHYRE2 protein structure prediction tool revealed altered conformational structures in both the Het and Hom lines. **C)** Western blotting confirmed reduced DUSP6 protein expression in the Het and complete absence in the Hom lines in a dosage-dependent manner. **D)** Immunocytochemical imaging of differentiated dopaminergic neurons derived from the iPSC lines shows a high purity of neurons expressing TH, MAP2 and DAPI nuclear stain.

The parental, Het, and Hom lines were differentiated into highly homogeneous dopaminergic neurons using an induction-based protocol (Powell et al., 2023). These neurons expressed TH, a key dopaminergic marker, at > 99% purity and also expressed the neuronal marker MAP2 (Figure 1D).

### *DUSP6* knockout reveals top DEGs are associated with dopaminergic signalling and neurodevelopment

To investigate transcriptional effects of *DUSP6* knockout, we performed whole-transcriptome RNA sequencing followed by pairwise differential expression analysis among parental, Het, and Hom lines. The parental and Hom comparison revealed 514 DEGs (|log2FC| ⩾ 2), with top 10 comprising extracellular matrix (ECM) and cell-adhesion molecules (*COL1A2, COL4A2, FBN1, TNC, CDH6*) or ECM associated/secreted neurodevelopmental regulators (*IGFBP7, FSTL1, SEMA3C, ANOS1, FKPB10*), all of which support neurogenesis, axon guidance, and synaptic plasticity (Figure 2A and Supplementary Table 3) (Bae et al., 2018; Chen et al., 2024; Hossain et al., 2025; Kumari et al., 2023; Lyons et al., 1994; Moreno-Flores et al., 2003; Nguyen et al., 2024; Osterhout et al., 2011; Steup et al., 2000; Takeuchi et al., 2015; Tucic et al., 2021).

**Figure 2.**
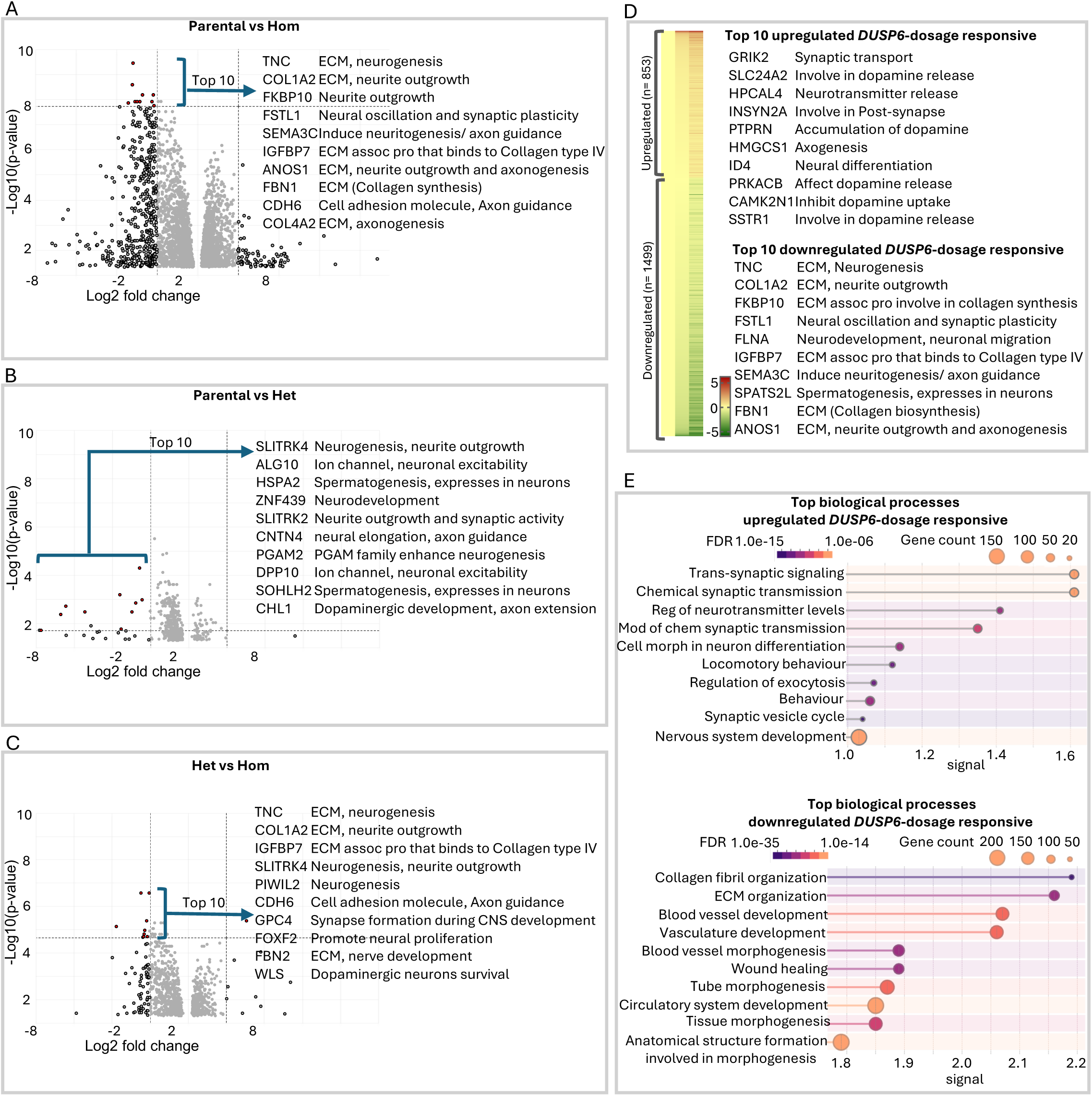
Differential gene expression analysis of the *DUSP6* knockout lines show high involvement in dopamine signalling and neurodevelopmental pathways. **A)** Volcano plot illustrating pairwise comparison of *DUSP6* knockout DEGs across parental vs Hom, **B)** Parental vs Het, and **C)** Het vs Hom lines. The top 10 high-effect-size DEGs (log2FC ⩾ 2), ranked by significance, are presented with their implicated neuronal function. **D)** Heat map of *DUSP6* dosage-responsive DEGs across parental, Het, and Hom lines, highlighting the top 10 up- and downregulated genes with their associated neuronal function. **E)** Gene enrichment analysis of dosage-responsive DEGs demonstrating that upregulated genes are predominantly associated with synaptic processes related to neurotransmitter signalling and neurogenesis related processes, whereas downregulated genes are enriched for general developmental and neurogenesis related pathways.

Comparisons of Het vs. Hom and parental vs. Het identified 77 and 25 DEGs (|log2FC| ⩾ 2) respectively (Figure 2AB–C, Supplementary Tables 4–5). The top 10 protein coding genes overlap functionally with those in the parental vs Hom comparison. These included genes involved in neurogenesis (*TNC, SLITRK4, PIWIL2, CPC4*, *ZNF439, SLITRK2, CNTN4, PGAM2, FOXF2, COL1A2, IGFBP7, CDH6,* and *FBN2*), several of which encode ECM associated molecules, as well as genes implicated in dopaminergic neurons survival and their pathway development (*WLS, CHL1*), and genes associated with neuronal excitability (*ALG10, DPP10*) (Al-Naama et al., 2020; Alsanie et al., 2017; Aruga & Mikoshiba, 2003; Bamford et al., 2024; Chen et al., 2014; Chernousov et al., 2010; El Chehadeh et al., 2022; Everson et al., 2017; Farhy-Tselnicker et al., 2017; Gasperini et al., 2023; Hoshi et al., 1998; Kim et al., 2020; Matsumoto et al., 2024; Prakash, 2022).

Across all comparisons, we further identified genes that altered in the same direction in both the Het and Hom lines (i.e., either both upregulated or both downregulated) with a greater fold change in the Hom line. These were defined as *DUSP6* dosage-responsive DEGs, with a total of 2,352 genes (1,499 downregulated and 853 upregulated) (Figure 2D, Supplementary Table 6). Among the top 10 upregulated genes, eight are involved in synaptic transmission and dopamine signalling (*GRIK2, SLC24A2, HPCAL4, INSYN2A, PTPRN, PRKACB, CAMK2N1 and SSTR1*); while the remaining two (*HMGCS1 and ID4*) are associated with neuronal development and differentiation (Bedford et al., 2005; Fog et al., 2006; Huang et al., 2019; Kershberg et al., 2022; Luderman et al., 2015; Mathews et al., 2014; Rajput et al., 2011; Sakharkar et al., 2019; Uezu et al., 2016; Wang et al., 2025). The enrichment of these dopamine related genes among the upregulated transcripts in both knockout lines aligns with previous evidence that *DUSP6* functions as a negative regulator of extracellular dopamine (Mortensen, 2013). The top downregulated genes are predominantly neurogenesis related and substantially overlap with those identified in the pairwise comparisons (*TNC, FSTL1, SEMA3C, FLNA, ANOS1, COL1A2, FKBP10, IGFBP7*, and *FBN1*) (Figure 2B). Across all comparisons, we also identified few spermatogenesis related genes (*SPATS2L HSPA2, SOHLH2*) with their neuronal function are currently unknown; however, prior studies have associated them with psychiatric disorders (Petyuk et al., 2018; Smeland et al., 2018).

Gene-enrichment analysis of the *DUSP6* dosage-responsive DEGs revealed that the upregulated genes are highly enriched for biological processes related to neurotransmitter synaptic transport (e.g. Chemical synaptic transmission (GO:0007268, FDR = 8.23 x 10^-15^), Neurotransmitter secretion (GO:0007269, FDR = 1.89 x 10^-5^)), and neurodevelopment (e.g. Neuron development (GO:0048666, FDR = 6.49 x 10^-10^), Axon development (GO:0061564, FDR = 1.04 x 10^-7^)), whereas the downregulated genes are predominantly associated with extracellular matrix associated and developmental processes (e.g. Extracellular matrix organization (GO:0030198, FDR = 3.66 x 10^-21^), Tube development (GO:0035295, FDR = 5.71 x 10^-34^)) (Figure 2E). Consistent with this, enrichment analysis of DEGs from the pairwise comparisons across the parental, Het, and Hom lines showed a similar pattern (Supplementary Figure 1–3). The convergence of shared functions across both pairwise comparisons and the dosage-responsive DEGs underscores the critical role of *DUSP6* in regulating dopaminergic signalling and neurodevelopment primarily through its dephosphorylation enzymatic activity (Seternes et al., 2019).

### *DUSP6* Knockout enhances dopamine secretion in a gene dosage-dependent manner independent of calcium channel alterations

To assess the hypothesis whether *DUSP6* knockout enhances dopamine secretion in ADHD dopaminergic neurons, we quantified both intracellular and extracellular dopamine across parental, Het, and Hom lines. Linear trend analysis revealed a significant increase in extracellular dopamine across the lines (Parental: Het: Hom; p = 0.015), with pairwise testing showing a significant elevation in the Hom line compared with the parental line (p = 0.016) (Figure 3A). These findings support a gene dosage-dependant increase in dopamine secretion following *DUSP6* knockout.

**Figure 3.**
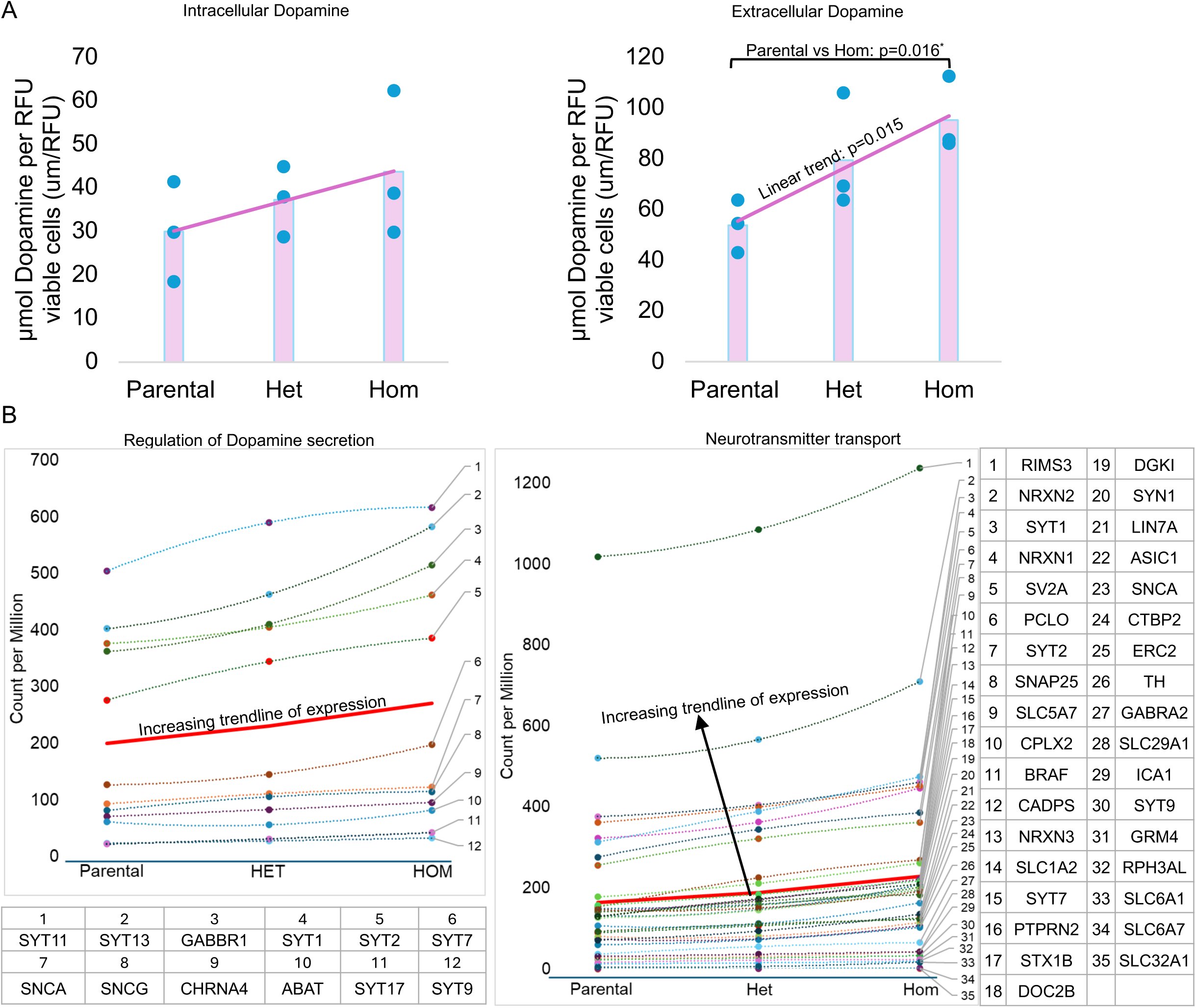
Dosage dependent regulatory effect of *DUSP6* on dopaminergic neurotransmission. **A)** A linear trend analysis shows a significant *DUSP6* dosage-dependent increase in extracellular dopamine across the Parental, Het and Hom line (p=0.015), with pairwise testing showing a significant elevation in the Hom line compared with the parental line (p = 0.016). The intracellular dopamine measurement did not show a significant change across the lines. Bars represent the mean of dopamine concentration for each group **B)** An increasing expression trend across parental, Het, and Hom lines was observed for DEGs associated with dopamine secretion (GO:0014059, FDR = 1.80 x 10^-5^) and neurotransmitter transport (GO:0006836, FDR = 1.3 x 10^-3^), which emerged as top enriched biological processes in the parental vs. Hom. The thick red line represents the trendline that best fits all data points.

This pattern aligns with transcriptional changes in neurotransmitter synaptic transport-related pathways that were enriched among the upregulated genes in *DUSP6* knockout lines. For instance, assessing DEGs from the two of the top dopamine-related biological processes including regulation of dopamine secretion (GO:0014059, FDR = 1.80 x 10^-5^) and neurotransmitter transport (GO:0006836, FDR = 1.3 x 10^-3^), has also revealed increasing expression across the parental, Het and Hom lines (Figure 3B).

Moreover, our data suggests that this elevation of extracellular dopamine is independent of CACNA1C-mediated calcium channel function. Contrary to our expectations, CACNA1C expression was not upregulated in the transcriptomic data following *DUSP6* knockout and calcium imaging showed no significant differences in either baseline or evoked calcium amplitudes among lines, indicating that *DUSP6* knockout does not markedly alter calcium signalling (Supplementary figure 4A–B). The frequency of spontaneously active cells similarly showed no significant variation (Supplementary figure 4C). These results suggest that the elevated extracellular dopamine is driven by upregulation of genes involved in synaptic machinery required for dopamine release, rather than by changes in calcium channel function.

### *DUSP6* knockout DEGs overlap with risk genes for other major psychiatric disorders

Given the association of *DUSP6* with multiple psychiatric conditions, we examined whether DEGs from the pairwise comparisons (parental, Het, and Hom) overlapped with GWAS-identified risk genes for major psychiatric disorders, including SCZ, MDD, BD, and ASD. DEGs from the parental vs. HOM comparison showed significant overlap with risk genes for BD (FDR = 7.0 × 10⁻⁵), MDD (FDR = 1.0 × 10⁻⁵), and SCZ (FDR = 1.0 × 10⁻⁵) (Figure 4A).

**Figure 4.**
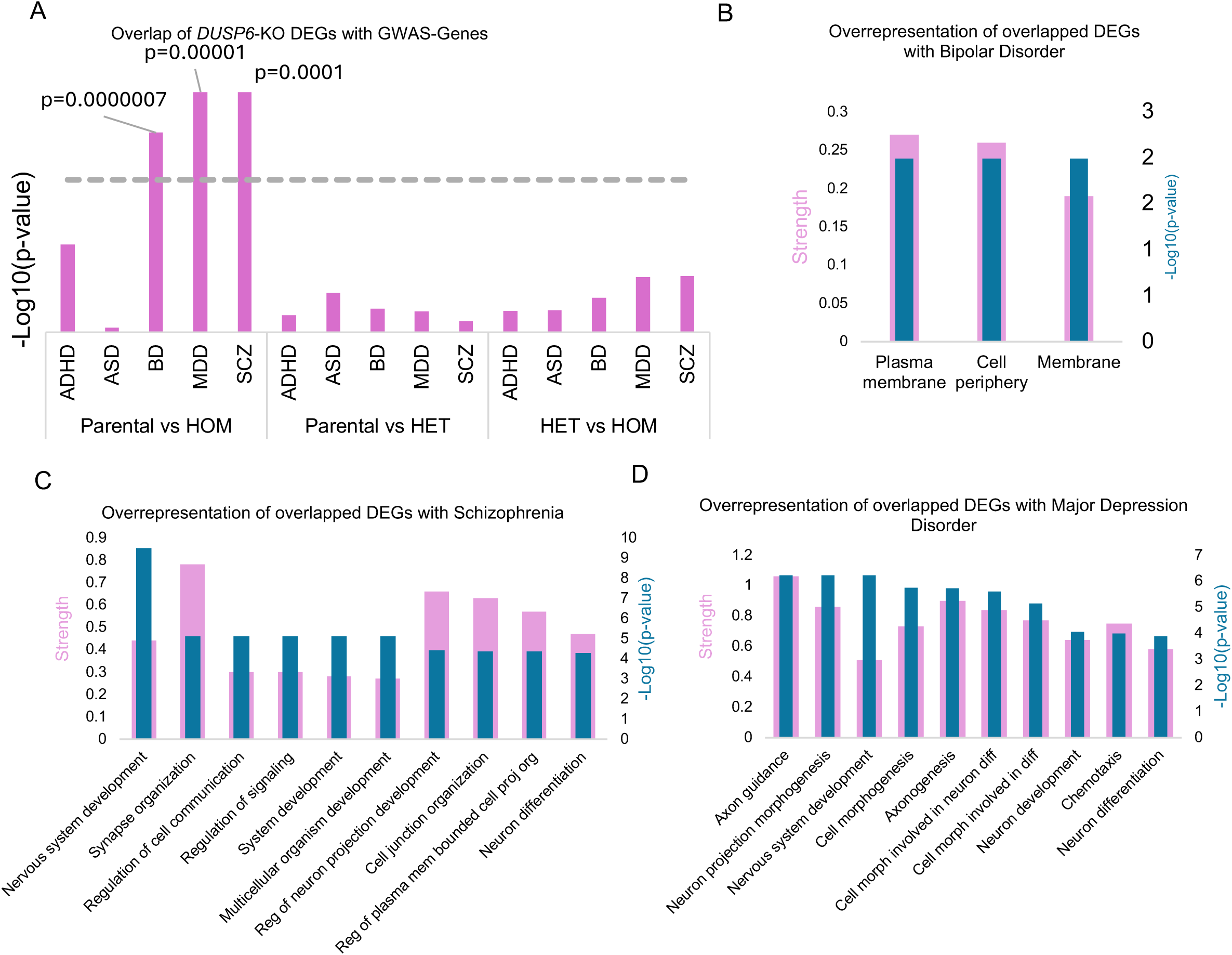
Overlap of *DUSP6*- knockout DEGs with GWAS-risk genes and enrichment analysis. **A)** *DUSP6* knockout DEGs show significant overlap with GWAS-identified risk genes for BD, MDD, and SCZ. **B)** Overrepresentation analysis of BP-overlapping DEGs reveals enrichment in biological processes, including membrane, plasma membrane and cell periphery. **C)** Top 10 biological processes, ranked by significance, identified through overrepresentation analysis of SCZ-overlapping DEGs. **D)** Top 10 biological processes, ranked by significance, identified through overrepresentation analysis of MDD-overlapping DEGs.

To further explore the biological significance of these overlaps, we performed enrichment analysis of overlapped DEGs for biological processes. The DEGs overlapping with BD risk genes were significantly overrepresented in processes related to the plasma membrane (GO:0005886) (FDR = 1.02 × 10⁻^2^), cell periphery (GO:0071944) (FDR = 1.02 × 10⁻^2^), and membrane GO:0071944 (FDR = 1.02 × 10⁻^2^) (Figure 4B and Supplementary Table 7). The DEGs overlapping with SCZ risk genes showed significant enrichment in biological processes associated with neurogenesis and synapse organization, such as nervous system development (GO:0007399, FDR = 1.02 × 10⁻^10^), synapse organization (GO:0050808, FDR = 7.51 × 10⁻^6^), and neuron differentiation (GO:0030182, FDR = 5.26 × 10⁻^5^) (Figure 4C and Supplementary Table 8). Similarly, the DEGs overlapping with MDD risk genes were significantly enriched in processes related to neurogenesis, axon guidance, and synapse assembly, such as axon guidance (GO:0007411, FDR = 3.99 × 10⁻^2^), synapse assembly (GO:0007416, FDR = 3.4 × 10⁻^14^), and axonogenesis (GO:0007409, FDR = 1.18 × 10⁻^2^) (Figure 4D and Supplementary Table 9). In contrast, DEGs from other pairwise comparisons, including parental versus Het and Het versus Hom, did not show nominal significance for overlap with risk genes of other psychiatric disorders after multiple correction.

## Discussion

In this study we highlight the potential role of the *DUSP6* gene in the pathophysiology of ADHD, specifically through its influence on dopaminergic neurotransmission. Our findings demonstrate a gene dosage-dependent regulatory effect of *DUSP6* on dopaminergic signalling, where knockout of the gene led to a linear upward increase in extracellular dopamine across the parental, heterozygous and homozygous lines. This pattern was consistent with the transcriptomic data at both pathway and gene level. We observed increasing expression of DEGs associated with top biological processes related to dopamine secretion and neurotransmitter transport. Further, most of the top upregulated *DUSP6*-dosage responsive DEGS appear to encode components or regulators of the synaptic machinery (e.g. *INSYN2A, PTPRN, SLC24A2, GRIK2, SSTR1*) effective for neurotransmitter release or act as regulators of dopaminergic neurotransmission (e.g. *HPCAL4, PRKACB, CAMK2N1*). This highlights the role of *DUSP6* in modulating the functional aspects of dopamine signalling at the synapse.

Our data indicate that *DUSP6* is not only a key regulator of dopaminergic neurotransmission but also an important modulator of neurodevelopment, representing another potential mechanism through which *DUSP6* may contribute to ADHD. Many of the upregulated genes showed enrichment for pathways involved in neurodevelopmental processes including axon development, neuron differentiation, and neurogenesis. In parallel, downregulated genes were predominantly enriched in pathways related to neuronal structure and developmental processes. Notably, a great portion of the top downregulated DEGs is related to extracellular matrix (ECM) and cell-adhesion molecules such as *COL1A2, TNC, FBN1, ANOS1, IGFBP7, CDH6*, all of which are implicated in neurogenesis related functions such as neurite outgrowth and axon guidance. Genes involved in collagen synthesis and assembly have previously been found to correlate significantly with ADHD in transcriptome-wide association studies (Demontis et al., 2021) and have been proposed as potential druggable targets for ADHD (Hegvik et al., 2021). In line with this, cadherin family genes has been linked to ADHD through multiple candidate and association studies (Hawi et al., 2018), further supporting the relevance of ECM and adhesion related pathways in ADHD biology.

The overlap of *DUSP6* knockout DEGs with those associated with other disorders, including major depression, bipolar disorder, and schizophrenia, suggests shared genetic pathways potentially driven by *DUSP6*. Downregulation of *DUSP6* expression has been reported in post-mortem brain tissue of patients with depressive disorders, as well as in those with bipolar disorder (Labonte et al., 2017; Struyf et al., 2008). Further, the increasing use of antipsychotic medications has been shown to downregulate *DUSP6* in the anterior cingulate cortex (ACC) of patients with schizophrenia, suggesting that *DUSP6* may have originally been upregulated in this disorder, and the antipsychotic medications may have targeted this gene, leading to its downregulation (Kondo et al., 2022). This indicates that both up- and down-regulation of *DUSP6* can lead to significant dysfunctions in key brain pathways, potentially contributing to the manifestation of these disorders.

Our findings suggests that *DUSP6* may have differing regulatory effects on calcium channels depending on whether it is upregulated or downregulated. While upregulation of *DUSP6* has been shown to downregulate *CACNA1C*, our findings indicate that the reverse is not necessarily true (Mortensen, 2013). This discrepancy could be attributed to previous work been conducted in a rat PC12 adrenal medullary tumor-derived cell line, whereas our experiments were performed in a human dopaminergic neuronal model. This highlights the importance of considering different cell types and species, as they may have distinct regulatory networks, underscoring the importance to study disorders in a cell-type-specific context. To gain a deeper understanding, follow-up experiments using CRISPR activation (CRISPRa) and CRISPR interference (CRISPRi) in more complex models, such as organoids, are recommended to better understand the effect of both over and under expression of *DUSP6*.

Our study, highlights the value of combining iPSC modelling with CRISPR technology, demonstrating that generating isogenic lines differing at only a single locus effectively overcomes the variability inherent in traditional case-control designs (Matos et al., 2020). Moreover, our findings emphasise the critical role of human models in psychiatric research, as they can capture disease-relevant biology that may not be fully recapitulated in animal models (Kampmann, 2020).

In conclusion, our study contributes to a deeper understanding of the molecular underpinnings of ADHD through the lens of GWAS identified risk gene in human neuronal model. Further, it underscores the importance of *DUSP6* as a regulator of both dopaminergic neurotransmission and neurodevelopment more generally. Given that GWAS risk genes typically exert small effect sizes through inherited causal SNPs that alter gene expression, future research should explore SNP level effects using CRISPRi/CRISPRa titration or CRISPR knock-in approaches. Additionally, investigating the downstream signalling pathways affected by *DUSP6* modulation will be essential for identifying potential targets for therapeutic interventions in ADHD.

## Acknowledgement

This work was supported by NHMRC grants: APP1146644 awarded to ZH and MAB, and APP2025415 and APP1154378 awarded to MAB, and the MIPS Neuroscience & Mental Health Therapeutic Program Area awarded to KJG & SH.

## References

Al-Naama, N., Mackeh, R., & Kino, T. (2020). C(2)H(2)-Type Zinc Finger Proteins in Brain Development, Neurodevelopmental, and Other Neuropsychiatric Disorders: Systematic Literature-Based Analysis. Front Neurol, 11, 32. 10.3389/fneur.2020.00032

Alsanie, W. F., Penna, V., Schachner, M., Thompson, L. H., & Parish, C. L. (2017). Homophilic binding of the neural cell adhesion molecule CHL1 regulates development of ventral midbrain dopaminergic pathways. Sci Rep, 7(1), 9368. 10.1038/s41598-017-09599-y

Aruga, J., & Mikoshiba, K. (2003). Identification and characterization of Slitrk, a novel neuronal transmembrane protein family controlling neurite outgrowth. Mol Cell Neurosci, 24(1), 117–129. 10.1016/s1044-7431(03)00129-5

Bae, C. J., Hong, C. S., & Saint-Jeannet, J. P. (2018). Anosmin-1 is essential for neural crest and cranial placodes formation in Xenopus. Biochem Biophys Res Commun, 495(3), 2257–2263. 10.1016/j.bbrc.2017.12.127

Bamford, R. A., Zuko, A., Eve, M., Sprengers, J. J., Post, H., Taggenbrock, R., Fabetaler, D., Mehr, A., Jones, O. J. R., Kudzinskas, A., Gandawijaya, J., Muller, U. C., Kas, M. J. H., Burbach, J. P. H., & Oguro-Ando, A. (2024). CNTN4 modulates neural elongation through interplay with APP. Open Biol, 14(5), 240018. 10.1098/rsob.240018

Bedford, L., Walker, R., Kondo, T., van Cruchten, I., King, E. R., & Sablitzky, F. (2005). Id4 is required for the correct timing of neural differentiation. Dev Biol, 280(2), 386–395. 10.1016/j.ydbio.2005.02.001

Chen, L., Hui, L., & Li, J. (2024). The multifaceted role of insulin-like growth factor binding protein 7. Front Cell Dev Biol, 12, 1420862. 10.3389/fcell.2024.1420862

Chen, T., Gai, W. P., & Abbott, C. A. (2014). Dipeptidyl peptidase 10 (DPP10(789)): a voltage gated potassium channel associated protein is abnormally expressed in Alzheimer’s and other neurodegenerative diseases. Biomed Res Int, 2014, 209398. 10.1155/2014/209398

Chernousov, M. A., Baylor, K., Stahl, R. C., Stecker, M. M., Sakai, L. Y., Lee-Arteaga, S., Ramirez, F., & Carey, D. J. (2010). Fibrillin-2 is dispensable for peripheral nerve development, myelination and regeneration. Matrix Biol, 29(5), 357–368. 10.1016/j.matbio.2010.02.006

de Leeuw, C. A., Mooij, J. M., Heskes, T., & Posthuma, D. (2015). MAGMA: generalized gene-set analysis of GWAS data. PLoS Comput Biol, 11(4), e1004219. 10.1371/journal.pcbi.1004219

Demontis, D., Walters, G. B., Athanasiadis, G., Walters, R., Therrien, K., Nielsen, T. T., Farajzadeh, L., Voloudakis, G., Bendl, J., Zeng, B., Zhang, W., Grove, J., Als, T. D., Duan, J., Satterstrom, F. K., Bybjerg-Grauholm, J., Baekved-Hansen, M., Gudmundsson, O. O., Magnusson, S. H.,…Borglum, A. D. (2023). Genome-wide analyses of ADHD identify 27 risk loci, refine the genetic architecture and implicate several cognitive domains. Nat Genet, 55(2), 198–208. 10.1038/s41588-022-01285-8

Demontis, D., Walters, R. K., Martin, J., Mattheisen, M., Als, T. D., Agerbo, E., Baldursson, G., Belliveau, R., Bybjerg-Grauholm, J., Baekvad-Hansen, M., Cerrato, F., Chambert, K., Churchhouse, C., Dumont, A., Eriksson, N., Gandal, M., Goldstein, J. I., Grasby, K. L., Grove, J.,…Neale, B. M. (2019). Discovery of the first genome-wide significant risk loci for attention deficit/hyperactivity disorder. Nat Genet, 51(1), 63–75. 10.1038/s41588-018-0269-7

Demontis, D., Walters, R. K., Rajagopal, V. M., Waldman, I. D., Grove, J., Als, T. D., Dalsgaard, S., Ribases, M., Bybjerg-Grauholm, J., Baekvad-Hansen, M., Werge, T., Nordentoft, M., Mors, O., Mortensen, P. B., Consortium, A. W. G. o. t. P. G., Cormand, B., Hougaard, D. M., Neale, B. M., Franke, B.,…Borglum, A. D. (2021). Risk variants and polygenic architecture of disruptive behavior disorders in the context of attention-deficit/hyperactivity disorder. Nat Commun, 12(1), 576. 10.1038/s41467-020-20443-2

Dobin, A., Davis, C. A., Schlesinger, F., Drenkow, J., Zaleski, C., Jha, S., Batut, P., Chaisson, M., & Gingeras, T. R. (2013). STAR: ultrafast universal RNA-seq aligner. Bioinformatics, 29(1), 15–21. 10.1093/bioinformatics/bts635

El Chehadeh, S., Han, K. A., Kim, D., Jang, G., Bakhtiari, S., Lim, D., Kim, H. Y., Kim, J., Kim, H., Wynn, J., Chung, W. K., Vitiello, G., Cutcutache, I., Page, M., Gecz, J., Harper, K., Han, A. R., Kim, H. M., Wessels, M., … Um, J. W. (2022). SLITRK2 variants associated with neurodevelopmental disorders impair excitatory synaptic function and cognition in mice. Nat Commun, 13(1), 4112. 10.1038/s41467-022-31566-z

Everson, J. L., Fink, D. M., Yoon, J. W., Leslie, E. J., Kietzman, H. W., Ansen-Wilson, L. J., Chung, H. M., Walterhouse, D. O., Marazita, M. L., & Lipinski, R. J. (2017). Sonic hedgehog regulation of Foxf2 promotes cranial neural crest mesenchyme proliferation and is disrupted in cleft lip morphogenesis. Development, 144(11), 2082–2091. 10.1242/dev.149930

Faraone, S. V., Bellgrove, M. A., Brikell, I., Cortese, S., Hartman, C. A., Hollis, C., Newcorn, J. H., Philipsen, A., Polanczyk, G. V., Rubia, K., Sibley, M. H., & Buitelaar, J. K. (2024). Attention-deficit/hyperactivity disorder. Nat Rev Dis Primers, 10(1), 11. 10.1038/s41572-024-00495-0

Farhy-Tselnicker, I., van Casteren, A. C. M., Lee, A., Chang, V. T., Aricescu, A. R., & Allen, N. J. (2017). Astrocyte-Secreted Glypican 4 Regulates Release of Neuronal Pentraxin 1 from Axons to Induce Functional Synapse Formation. Neuron, 96(2), 428–445 e413. 10.1016/j.neuron.2017.09.053

Felix Krueger, F. J., Phil Ewels, Ebrahim Afyounian, & Benjamin Schuster-Boeckler.. (2021). FelixKrueger/TrimGalore: v0.6.7 - DOI via Zenodo (0.6.7). Zenodo. 10.5281/zenodo.5127899

Fog, J. U., Khoshbouei, H., Holy, M., Owens, W. A., Vaegter, C. B., Sen, N., Nikandrova, Y., Bowton, E., McMahon, D. G., Colbran, R. J., Daws, L. C., Sitte, H. H., Javitch, J. A., Galli, A., & Gether, U. (2006). Calmodulin kinase II interacts with the dopamine transporter C terminus to regulate amphetamine-induced reverse transport. Neuron, 51(4), 417–429. 10.1016/j.neuron.2006.06.028

Gasperini, C., Tuntevski, K., Beatini, S., Pelizzoli, R., Lo Van, A., Mangoni, D., Cossu, R. M., Pascarella, G., Bianchini, P., Bielefeld, P., Scarpato, M., Pons-Espinal, M., Sanges, R., Diaspro, A., Fitzsimons, C. P., Carninci, P., Gustincich, S., & De Pietri Tonelli, D. (2023). Piwil2 (Mili) sustains neurogenesis and prevents cellular senescence in the postnatal hippocampus. EMBO Rep, 24(2), e53801. 10.15252/embr.202153801

Harshil Patel, P. E., Alexander Peltzer, Olga Botvinnik, Gregor Sturm, Denis Moreno, Pranathi Vemuri, silviamorins, Lorena Pantano, Mahesh Binzer-Panchal, nf-core bot, Gavin Kelly, Maxime U. Garcia, FriederikeHanssen, Matthias Zepper, James A. Fellows Yates, Chris Cheshire, rfenouil, Jose Espinosa-Carrasco, … George Hall. (2023). nf-core/rnaseq: nf-core/rnaseq v3.10.1 - Plastered Rhodium Rudolph (3.10.1). Zenodo. 10.5281/zenodo.7505987

Hawi, Z., Tong, J., Dark, C., Yates, H., Johnson, B., & Bellgrove, M. A. (2018). The role of cadherin genes in five major psychiatric disorders: A literature update. Am J Med Genet B Neuropsychiatr Genet, 177(2), 168–180. 10.1002/ajmg.b.32592

Hegvik, T. A., Waloen, K., Pandey, S. K., Faraone, S. V., Haavik, J., & Zayats, T. (2021). Druggable genome in attention deficit/hyperactivity disorder and its co-morbid conditions. New avenues for treatment. Mol Psychiatry, 26(8), 4004–4015. 10.1038/s41380-019-0540-z

Hoshi, N., Takahashi, H., Shahidullah, M., Yokoyama, S., & Higashida, H. (1998). KCR1, a membrane protein that facilitates functional expression of non-inactivating K+ currents associates with rat EAG voltage-dependent K+ channels. J Biol Chem, 273(36), 23080–23085. 10.1074/jbc.273.36.23080

Hossain, A. S., Clarin, M., Kimura, K., Biggin, G., Taga, Y., Uto, K., Yamagishi, A., Motoyama, E., Narenmandula, Mizuno, K., Nakamura, C., Asano, K., Ohtsuki, S., Nakamura, T., Kanki, S., Baldock, C., Raja, E., & Yanagisawa, H. (2025). Fibrillin-1 G234D mutation in the hybrid1 domain causes tight skin associated with dysregulated elastogenesis and increased collagen cross-linking in mice. Matrix Biol, 135, 24–38. 10.1016/j.matbio.2024.11.006

Huang, X., Wang, M., Zhang, Q., Chen, X., & Wu, J. (2019). The role of glutamate receptors in attention-deficit/hyperactivity disorder: From physiology to disease. Am J Med Genet B Neuropsychiatr Genet, 180(4), 272–286. 10.1002/ajmg.b.32726

Kampmann, M. (2020). CRISPR-based functional genomics for neurological disease. Nat Rev Neurol, 16(9), 465–480. 10.1038/s41582-020-0373-z

Kelley, L. A., Mezulis, S., Yates, C. M., Wass, M. N., & Sternberg, M. J. (2015). The Phyre2 web portal for protein modeling, prediction and analysis. Nat Protoc, 10(6), 845–858. 10.1038/nprot.2015.053

Kershberg, L., Banerjee, A., & Kaeser, P. S. (2022). Protein composition of axonal dopamine release sites in the striatum. Elife, 11. 10.7554/eLife.83018

Kim, W., Kwon, H. J., Jung, H. Y., Yoo, D. Y., Kim, D. W., & Hwang, I. K. (2020). Phosphoglycerate mutase 1 reduces neuronal damage in the hippocampus following ischemia/reperfusion through the facilitation of energy utilization. Neurochem Int, 133, 104631. 10.1016/j.neuint.2019.104631

Kondo, M. A., Norris, A. L., Yang, K., Cheshire, M., Newkirk, I., Chen, X., Ishizuka, K., Jaffe, A. E., Sawa, A., & Pevsner, J. (2022). Dysfunction of mitochondria and GABAergic interneurons in the anterior cingulate cortex of individuals with schizophrenia. Neurosci Res, 185, 67–72. 10.1016/j.neures.2022.09.011

Kumari, E., Xu, A., Chen, R., Yan, Y., Yang, Z., & Zhang, T. (2023). FSTL1-knockdown improves neural oscillation via decreasing neuronal-inflammation regulating apoptosis in Abeta(1-42) induced AD model mice. Exp Neurol, 359, 114231. 10.1016/j.expneurol.2022.114231

Labonte, B., Engmann, O., Purushothaman, I., Menard, C., Wang, J., Tan, C., Scarpa, J. R., Moy, G., Loh, Y. E., Cahill, M., Lorsch, Z. S., Hamilton, P. J., Calipari, E. S., Hodes, G. E., Issler, O., Kronman, H., Pfau, M., Obradovic, A. L. J., Dong, Y.,…Nestler, E. J. (2017). Sex-specific transcriptional signatures in human depression. Nat Med, 23(9), 1102–1111. 10.1038/nm.4386

Law, C. W., Chen, Y., Shi, W., & Smyth, G. K. (2014). voom: Precision weights unlock linear model analysis tools for RNA-seq read counts. Genome Biol, 15(2), R29. 10.1186/gb-2014-15-2-r29

Luderman, K. D., Chen, R., Ferris, M. J., Jones, S. R., & Gnegy, M. E. (2015). Protein kinase C beta regulates the D(2)-like dopamine autoreceptor. Neuropharmacology, 89, 335–341. 10.1016/j.neuropharm.2014.10.012

Lyons, W. E., George, E. B., Dawson, T. M., Steiner, J. P., & Snyder, S. H. (1994). Immunosuppressant FK506 promotes neurite outgrowth in cultures of PC12 cells and sensory ganglia. Proc Natl Acad Sci U S A, 91(8), 3191–3195. 10.1073/pnas.91.8.3191

Mathews, E. S., Mawdsley, D. J., Walker, M., Hines, J. H., Pozzoli, M., & Appel, B. (2014). Mutation of 3-hydroxy-3-methylglutaryl CoA synthase I reveals requirements for isoprenoid and cholesterol synthesis in oligodendrocyte migration arrest, axon wrapping, and myelin gene expression. J Neurosci, 34(9), 3402–3412. 10.1523/JNEUROSCI.4587-13.2014

Matos, M. R., Ho, S. M., Schrode, N., & Brennand, K. J. (2020). Integration of CRISPR-engineering and hiPSC-based models of psychiatric genomics. Mol Cell Neurosci, 107, 103532. 10.1016/j.mcn.2020.103532

Matsumoto, Y., Miwa, H., Katayama, K. I., Watanabe, A., Yamada, K., Ito, T., Nakagawa, S., & Aruga, J. (2024). Slitrk4 is required for the development of inhibitory neurons in the fear memory circuit of the lateral amygdala. Front Mol Neurosci, 17, 1386924. 10.3389/fnmol.2024.1386924

Moreno-Flores, M. T., Martin-Aparicio, E., Martin-Bermejo, M. J., Agudo, M., McMahon, S., Avila, J., Diaz-Nido, J., & Wandosell, F. (2003). Semaphorin 3C preserves survival and induces neuritogenesis of cerebellar granule neurons in culture. J Neurochem, 87(4), 879–890. 10.1046/j.1471-4159.2003.02051.x

Mortensen, O. V. (2013). MKP3 eliminates depolarization-dependent neurotransmitter release through downregulation of L-type calcium channel Cav1.2 expression. Cell Calcium, 53(3), 224–230. 10.1016/j.ceca.2012.12.004

Mortensen, O. V., Larsen, M. B., Prasad, B. M., & Amara, S. G. (2008). Genetic complementation screen identifies a mitogen-activated protein kinase phosphatase, MKP3, as a regulator of dopamine transporter trafficking. Mol Biol Cell, 19(7), 2818-2829. 10.1091/mbc.e07-09-0980

Namipashaki, A., Pugsley, K., Liu, X., Abrehart, K., Lim, S. M., Sun, G., Herold, M. J., Polo, J. M., Bellgrove, M. A., & Hawi, Z. (2023). Integration of xeno-free single-cell cloning in CRISPR-mediated DNA editing of human iPSCs improves homogeneity and methodological efficiency of cellular disease modeling. Stem Cell Reports, 18(12), 2515–2527. 10.1016/j.stemcr.2023.10.013

Nguyen, U. N., Lee, F. S., Caparaso, S. M., Leoni, J. T., Redwine, A. L., & Wachs, R. A. (2024). Type I collagen concentration affects neurite outgrowth of adult rat DRG explants by altering mechanical properties of hydrogels. J Biomater Sci Polym Ed, 35(2), 164–189. 10.1080/09205063.2023.2272479

Osterhout, J. A., Josten, N., Yamada, J., Pan, F., Wu, S. W., Nguyen, P. L., Panagiotakos, G., Inoue, Y. U., Egusa, S. F., Volgyi, B., Inoue, T., Bloomfield, S. A., Barres, B. A., Berson, D. M., Feldheim, D. A., & Huberman, A. D. (2011). Cadherin-6 mediates axon-target matching in a non-image-forming visual circuit. Neuron, 71(4), 632–639. 10.1016/j.neuron.2011.07.006

Patro, R., Duggal, G., Love, M. I., Irizarry, R. A., & Kingsford, C. (2017). Salmon provides fast and bias-aware quantification of transcript expression. Nat Methods, 14(4), 417–419. 10.1038/nmeth.4197

Petyuk, V. A., Chang, R., Ramirez-Restrepo, M., Beckmann, N. D., Henrion, M. Y. R., Piehowski, P. D., Zhu, K., Wang, S., Clarke, J., Huentelman, M. J., Xie, F., Andreev, V., Engel, A., Guettoche, T., Navarro, L., De Jager, P., Schneider, J. A., Morris, C. M., McKeith, I. G.,…Myers, A. J. (2018). The human brainome: network analysis identifies HSPA2 as a novel Alzheimer’s disease target. Brain, 141(9), 2721–2739. 10.1093/brain/awy215

Powell, S. K., O’Shea, C., Townsley, K., Prytkova, I., Dobrindt, K., Elahi, R., Iskhakova, M., Lambert, T., Valada, A., Liao, W., Ho, S. M., Slesinger, P. A., Huckins, L. M., Akbarian, S., & Brennand, K. J. (2023). Induction of dopaminergic neurons for neuronal subtype-specific modeling of psychiatric disease risk. Mol Psychiatry, 28(5), 1970–1982. 10.1038/s41380-021-01273-0

Powell., D. (2019). drpowell/degust 4.1.1 (4.1.1). Zenodo. 10.5281/zenodo.3501067

Prakash, N. (2022). Developmental pathways linked to the vulnerability of adult midbrain dopaminergic neurons to neurodegeneration. Front Mol Neurosci, 15, 1071731. 10.3389/fnmol.2022.1071731

Rajput, P. S., Kharmate, G., Norman, M., Liu, S. H., Sastry, B. R., Brunicardi, C. F., & Kumar, U. (2011). Somatostatin receptor 1 and 5 double knockout mice mimic neurochemical changes of Huntington’s disease transgenic mice. PLoS One, 6(9), e24467. 10.1371/journal.pone.0024467

Sakharkar, M. K., Kashmir Singh, S. K., Rajamanickam, K., Mohamed Essa, M., Yang, J., & Chidambaram, S. B. (2019). A systems biology approach towards the identification of candidate therapeutic genes and potential biomarkers for Parkinson’s disease. PLoS One, 14(9), e0220995. 10.1371/journal.pone.0220995

Seternes, O. M., Kidger, A. M., & Keyse, S. M. (2019). Dual-specificity MAP kinase phosphatases in health and disease. Biochim Biophys Acta Mol Cell Res, 1866(1), 124–143. 10.1016/j.bbamcr.2018.09.002

Smeland, O. B., Wang, Y., Frei, O., Li, W., Hibar, D. P., Franke, B., Bettella, F., Witoelar, A., Djurovic, S., Chen, C. H., Thompson, P. M., Dale, A. M., & Andreassen, O. A. (2018). Genetic Overlap Between Schizophrenia and Volumes of Hippocampus, Putamen, and Intracranial Volume Indicates Shared Molecular Genetic Mechanisms. Schizophr Bull, 44(4), 854–864. 10.1093/schbul/sbx148

Soneson, C., Love, M. I., & Robinson, M. D. (2015). Differential analyses for RNA-seq: transcript-level estimates improve gene-level inferences. F1000Res, 4, 1521. 10.12688/f1000research.7563.2

Steup, A., Lohrum, M., Hamscho, N., Savaskan, N. E., Ninnemann, O., Nitsch, R., Fujisawa, H., Puschel, A. W., & Skutella, T. (2000). Sema3C and netrin-1 differentially affect axon growth in the hippocampal formation. Mol Cell Neurosci, 15(2), 141–155. 10.1006/mcne.1999.0818

Struyf, J., Dobrin, S., & Page, D. (2008). Combining gene expression, demographic and clinical data in modeling disease: a case study of bipolar disorder and schizophrenia. BMC Genomics, 9, 531. 10.1186/1471-2164-9-531

Szklarczyk, D., Gable, A. L., Nastou, K. C., Lyon, D., Kirsch, R., Pyysalo, S., Doncheva, N. T., Legeay, M., Fang, T., Bork, P., Jensen, L. J., & von Mering, C. (2021). The STRING database in 2021: customizable protein-protein networks, and functional characterization of user-uploaded gene/measurement sets. Nucleic Acids Res, 49(D1), D605–D612. 10.1093/nar/gkaa1074

Szklarczyk, D., Kirsch, R., Koutrouli, M., Nastou, K., Mehryary, F., Hachilif, R., Gable, A. L., Fang, T., Doncheva, N. T., Pyysalo, S., Bork, P., Jensen, L. J., & von Mering, C. (2023). The STRING database in 2023: protein-protein association networks and functional enrichment analyses for any sequenced genome of interest. Nucleic Acids Res, 51(D1), D638–D646. 10.1093/nar/gkac1000

Takeuchi, M., Yamaguchi, S., Yonemura, S., Kakiguchi, K., Sato, Y., Higashiyama, T., Shimizu, T., & Hibi, M. (2015). Type IV Collagen Controls the Axogenesis of Cerebellar Granule Cells by Regulating Basement Membrane Integrity in Zebrafish. PLoS Genet, 11(10), e1005587. 10.1371/journal.pgen.1005587

Tong, J., Lee, K. M., Liu, X., Nefzger, C. M., Vijayakumar, P., Hawi, Z., Pang, K. C., Parish, C. L., Polo, J. M., & Bellgrove, M. A. (2019). Generation of four iPSC lines from peripheral blood mononuclear cells (PBMCs) of an attention deficit hyperactivity disorder (ADHD) individual and a healthy sibling in an Australia-Caucasian family. Stem Cell Res, 34, 101353. 10.1016/j.scr.2018.11.014

Tucic, M., Stamenkovic, V., & Andjus, P. (2021). The Extracellular Matrix Glycoprotein Tenascin C and Adult Neurogenesis. Front Cell Dev Biol, 9, 674199. 10.3389/fcell.2021.674199

Uezu, A., Kanak, D. J., Bradshaw, T. W., Soderblom, E. J., Catavero, C. M., Burette, A. C., Weinberg, R. J., & Soderling, S. H. (2016). Identification of an elaborate complex mediating postsynaptic inhibition. Science, 353(6304), 1123–1129. 10.1126/science.aag0821

Wang, Y. M., Wang, W. C., Pan, Y., Zeng, L., Wu, J., Wang, Z. B., Zhuang, X. L., Li, M. L., Cooper, D. N., Wang, S., Shao, Y., Wang, L. M., Fan, Y. Y., He, Y., Hu, X. T., & Wu, D. D. (2025). Regional and aging-specific cellular architecture of non-human primate brains. Genome Med, 17(1), 41. 10.1186/s13073-025-01469-x

