## supplementary figures for "*DUSP6* loss enhances dopamine secretion and modulate neurodevelopment in ADHD iPSC-derived dopaminergic neurons"

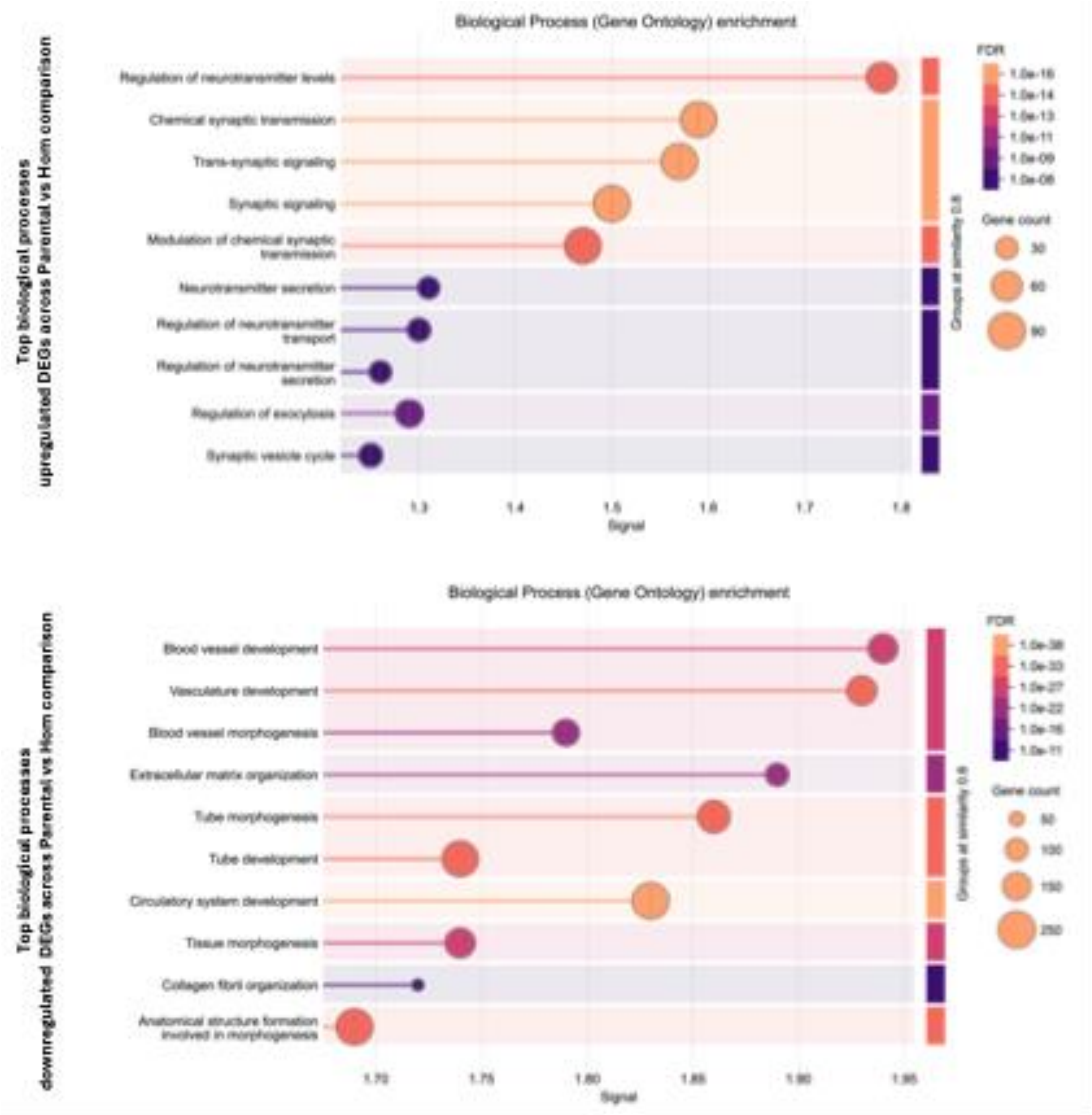


**Supplementary figure 1. Top biological processes enriched for the DEGs across Parental vs Hom comparison.**

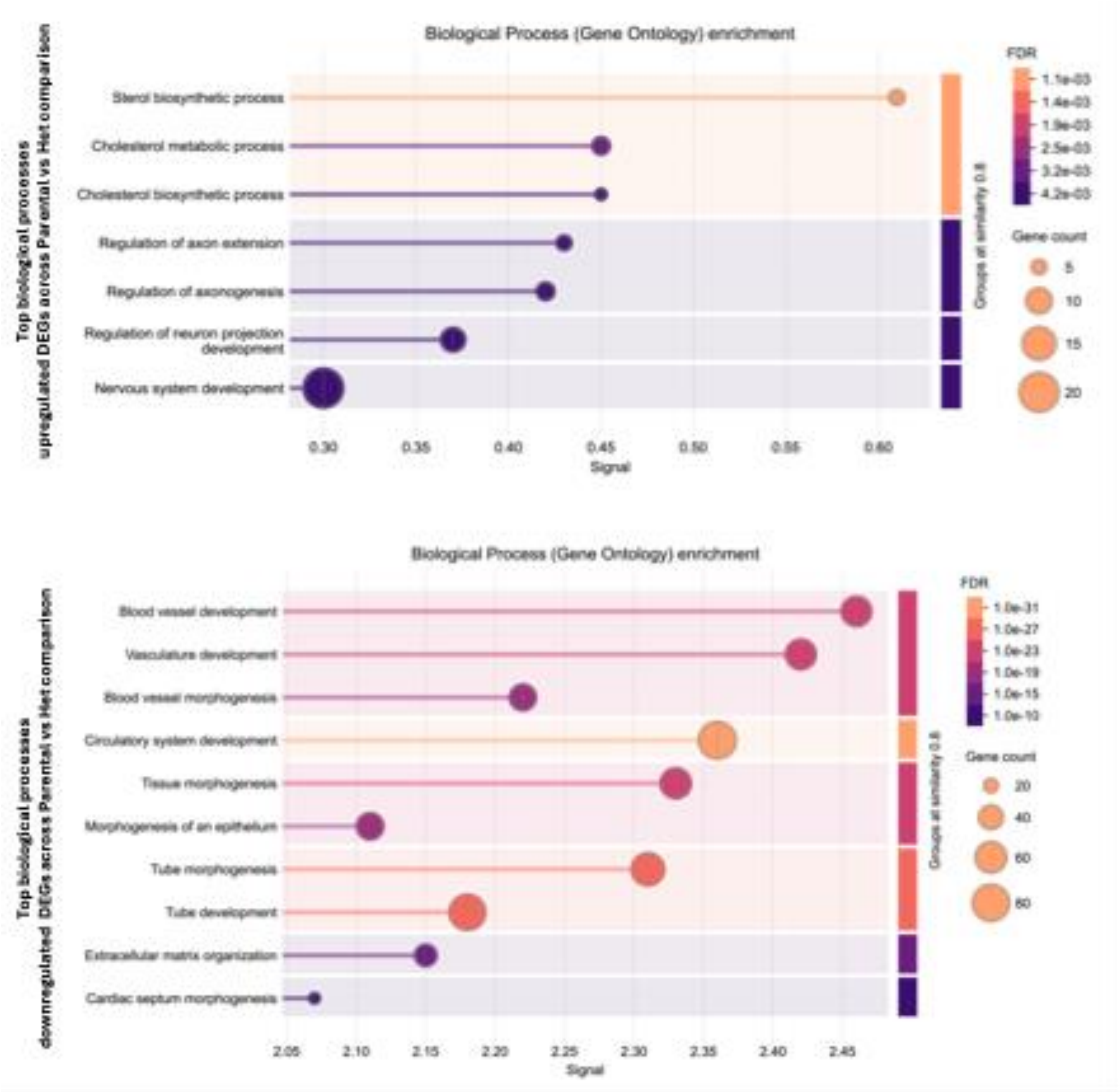


**Supplementary figure2. Top biological processes enriched for the DEGs across Parental vs Het comparison.**

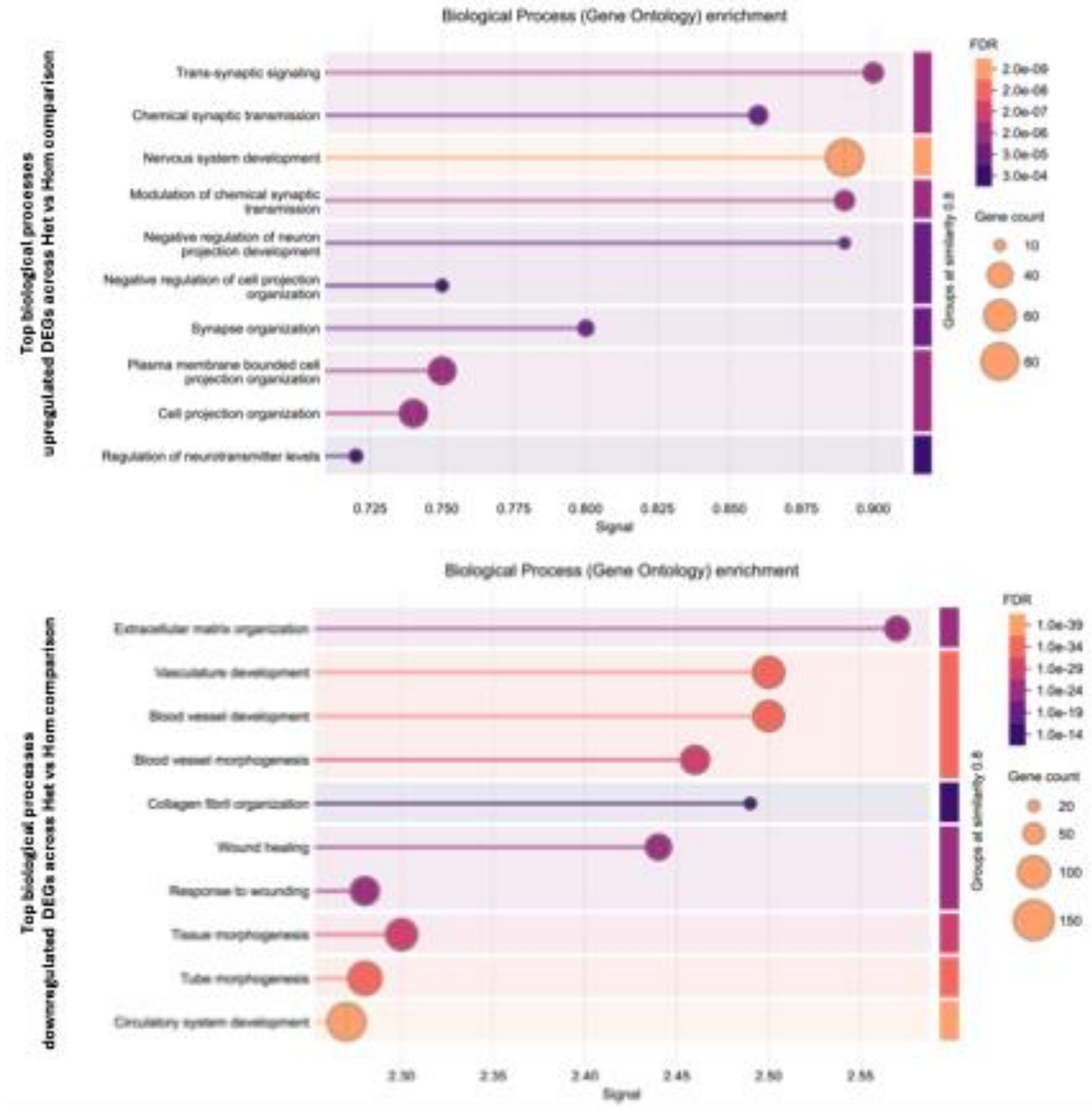


**Supplementary figure 3. Top biological processes enriched for the DEGs across Het vs Hom comparison.**

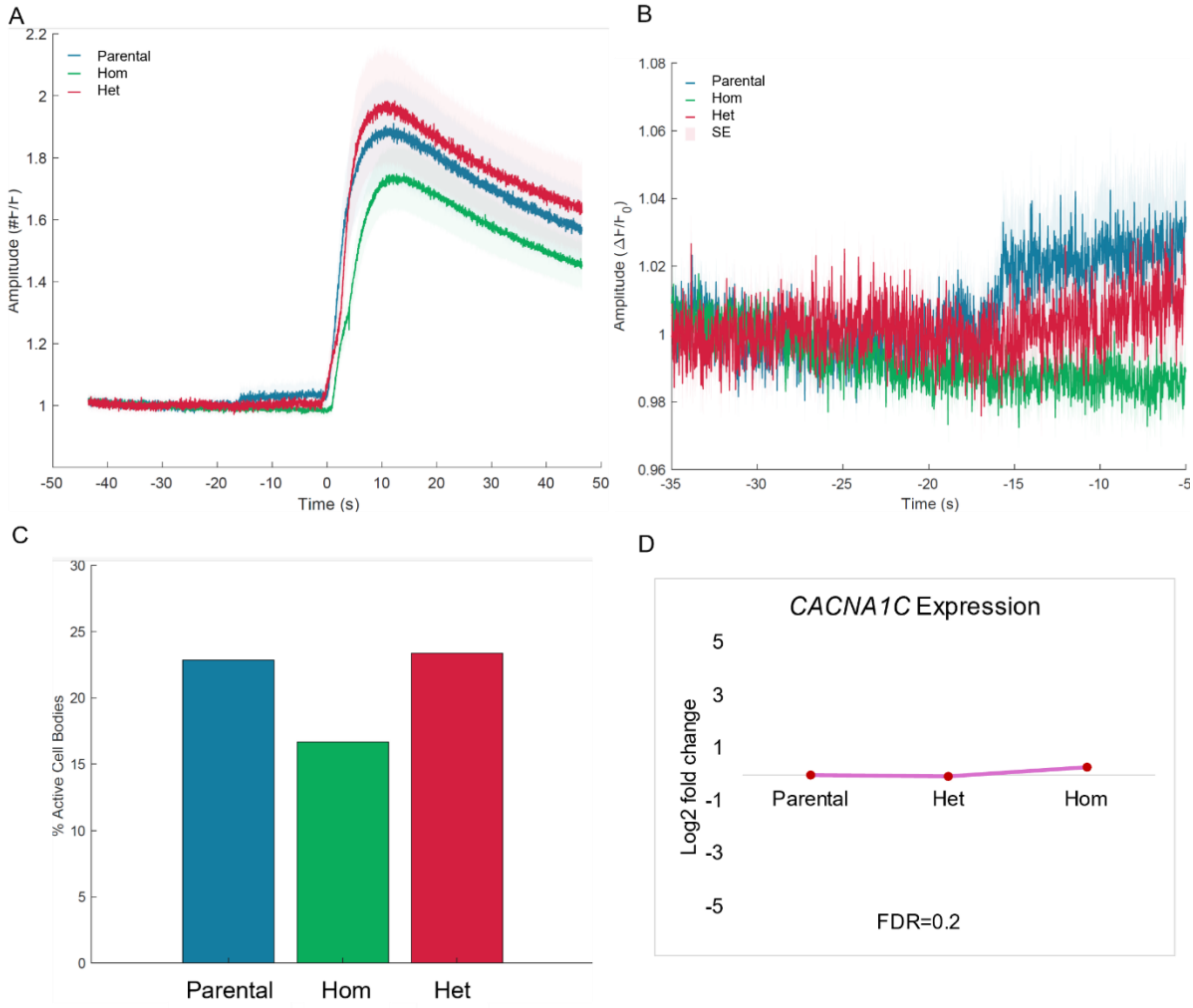


**Supplementary figure 4. Effect of *DUSP6* knockout on *CACNA1C* calcium channel in dopaminergic neurons.** Measurements of **A**) evoked and **B)** baseline amplitude show no significant differences across the parental, Het, and Hom lines. **C)** Similarly, no significant differences were observed in the frequency of spontaneously active cells across the lines. **D)** Consistently, RNA-sequencing results showed *CACNA1C* expression is not significantly altered due to the knockout of *DUSP6* in either the Het or Hom lines.
